# Chromosome dynamics and spatial organization during the non-binary cell cycle of a predatory bacterium

**DOI:** 10.1101/2020.12.21.419010

**Authors:** Jovana Kaljević, Terrens N. V. Saaki, Sander K. Govers, Ophélie Remy, Renske van Raaphorst, Thomas Lamot, Géraldine Laloux

**Affiliations:** de Duve Institute, UCLouvain, Brussels, Belgium; Howard Hughes Medical Institute, Stanford University, Stanford, CA 94305, USA

**Author notes:** equal contribution (Jovana Kaljević, Terrens N. V. Saaki).

## Abstract

In bacteria, the dynamics of chromosome replication and segregation are tightly coordinated with cell cycle progression, and largely rely on specific spatiotemporal arrangement of the chromosome. Whereas these key processes are mostly investigated in species that divide by binary fission, they remain mysterious in bacteria producing larger number of descendants. Here, we establish the predatory bacterium *Bdellovibrio bacteriovorus* as a model to investigate the non-binary processing of a circular chromosome. Our data reveal its extreme compaction in a dense polarized nucleoid. We also show that a first binary-like cycle of replication and asymmetric segregation is followed by multiple asynchronous rounds of replication and progressive ParAB*S*-dependent partitioning, uncoupled from cell division. Surprisingly, ParB localization at the centromere is cell-cycle regulated. Altogether, our findings support a model of complex chromosome choreography, leading to the generation of variable numbers of offspring, highlighting the adaptation of conserved mechanisms to achieve non-binary reproduction in bacteria.

## Introduction

Bacteria thrive in highly diverse environments to which they finely adapt, as illustrated by the immense variety of proliferation modes that were selected through evolution. Despite this tremendous diversity, most of our knowledge about bacterial multiplication derives from work on a subset of model species, which all divide by binary fission: one mother cell elongates, duplicates its genetic information and gives rise to two daughter cells upon a single cell division event (1). However, not all bacteria adhere to the simple paradigm of binary reproduction (2). Species from various lineages (including Actinobacteria, Cyanobacteria and Bdellovibrionata) rely on sophisticated cell cycles involving multiple fission (3–5). Here, larger and sometimes variable numbers of progeny are generated from a polyploid mother cell, through several (sequential or synchronous) septation events. The non-binary proliferation of these species, which is inherently complex due to the production of more than two descendants, offers an attractive platform to shed light on overlooked cell cycle regulation strategies in bacteria.

Even the seemingly simple cell cycles must be achieved with precision. For this, bacterial cells rely on specific and elaborate spatiotemporal organization to orchestrate key cellular processes. A prominent example of cellular organization in bacteria is the intricate coordination, in both space and time, of chromosome replication and segregation with other cell cycle events, including growth and cell division (6). In model bacteria, the spatial and temporal organization of the chromosome depends on specific subcellular positions of key chromosomal loci, mainly the replication origin (*ori*) and terminus (*ter*), which display highly regulated dynamics during the cell cycle (7–17). Soon after replication initiation, most species studied so far (with the exception of γ-proteobacteria) employ the ParAB*S* system to actively partition sister *ori* (18). In this system, directionality of *ori* segregation is provided by the exquisite interplay between the ParB protein, which binds and spreads from the centromeric *parS* sites near *ori*, and the unspecific DNA-binding ATPase ParA. Iterations of ParB-triggered ATPase activity and dissociation of ParA from the chromosome result in the segregation of duplicated ParB•*parS* complexes, before the replication forks reach the chromosomal *ter* (19,20). Specific mechanisms physically connect the partitioning of *ter* copies with cell constriction, thereby coordinating the last steps of chromosome segregation and cell division (21–23) and ensuring that each daughter cell is equipped with a full set of genetic material. Except in some streptomycetes (24,25), the spatiotemporal organization of the chromosome and the interplay between fundamental cellular processes remain essentially unexplored in non-binary growing bacteria.

The Gram-negative bacterium *B. bacteriovorus* (a member of the recently proposed Bdellovibrionata phylum (26), formerly considered as a δ-proteobacterium) features an extraordinary non-binary cell cycle (reviewed in (27)) (Figure 1). First, the mono-flagellated and fast-swimming attack phase (AP) cells search for prey in the environment. Upon pili-mediated attachment to a Gram-negative bacterium (28), the predator invades the periplasm of its prey by sneaking through a narrow hole in the outer membrane and peptidoglycan (29,30). Precise remodeling of the prey cell wall leads to rounding of the bdelloplast (*i.e.* the *Bdellovibrio-*infected prey bacterium) without altering its osmotic integrity (29,30). Within the bdelloplast, *B. bacteriovorus* digests the prey content (31) and grows as a filament before producing a variable, even or odd number of daughter cells, by multiple synchronous division events (32). Flagellated monoploid cells then exit the prey ghost (33) and resume the cycle.

**Figure 1.**
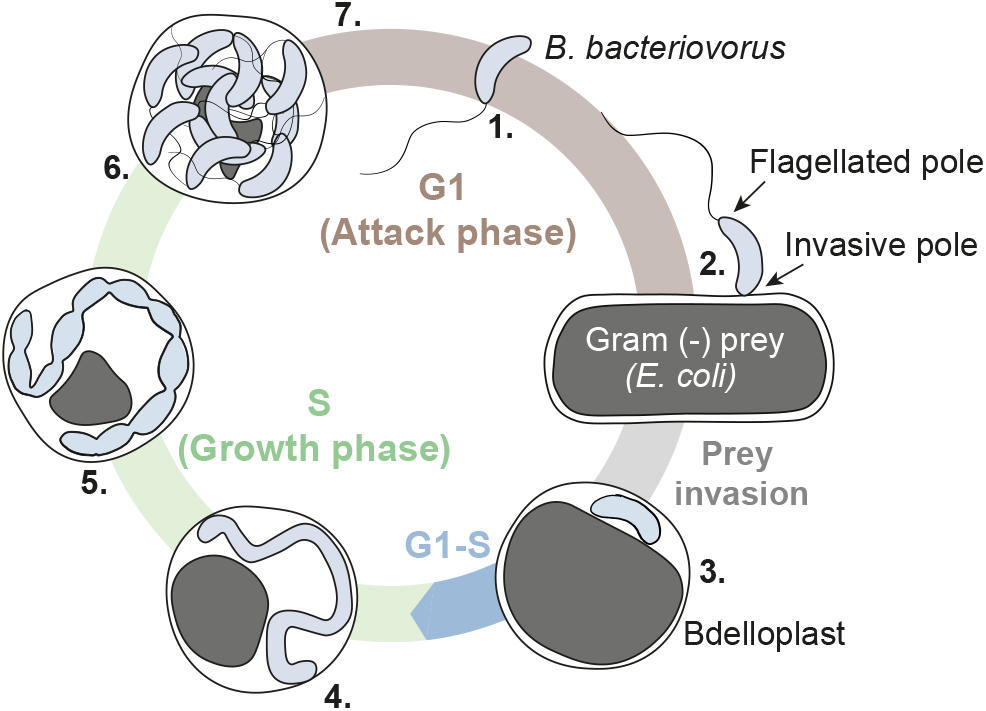
Schematics of the *Bdellovibrio bacteriovorus* cell cycle. Numbers indicate key steps in the cycle: **1.** Freely-swimming attack phase (AP) cells, **2.** Attachment of *B. bacteriovorus* to its prey, via pili located at the non-flagellated (invasive) pole, **3.** *B. bacteriovorus* resides in the periplasm of the prey, which is now called bdelloplast, **4.** Filamentous growth and consumption of prey content, **5.** Pre-divisional state, **6.** Non-binary division of the mother cell generates an odd or even number of daughter cells, which mature before **7.** escaping the prey remnants and resuming the cell cycle. Attack phase (AP) and growth phase (GP) represent the G1 (non-replicative) and S (replicative) stages of the cell cycle, respectively. A G1-S transition takes place upon prey invasion (see results).

The filamentous growth and multiple progeny of *B. bacteriovorus* raise unexplored fundamental questions regarding the orchestration of chromosome-related processes during the cell cycle (34). After prey invasion, the single circular chromosome of each *B. bacteriovorus* cell (35) must be copied and partitioned multiple times before releasing daughter cells, challenging the principle of temporal coupling between chromosome replication, segregation, and cell division (36). Remarkably, chromosome replication must be tuned such that odd or even numbers can be obtained, in contrast with common exponential multiplication patterns. Whereas a few hints suggest that *B. bacteriovorus* exploits multiple replisomes at the same time during its growth phase (37,38), unambiguous insights into native replication dynamics are lacking. In addition, how and when chromosome segregation occurs relative to replication and cell division is unknown.

Here we set out to provide key insights into the spatial organization of the chromosome and shed light on the orchestration of chromosome replication and segregation during the intriguing lifecycle of *B. bacteriovorus*. Using epifluorescence microscopy on living cells, followed by quantitative image analysis at the single-cell and population levels, we monitored the subcellular localization of *ori* and *ter* loci, as well as the native replication and segregation machineries, during the G1 (non-replicative), G1-S transition and S (replicative) phases of the synchronized predatory cell cycle. Our results reveal the longitudinal organization of the chromosome in *B. bacteriovorus*, its unique polarity and compaction, and allow us to propose a model for the complex choreography of chromosome replication and ParAB*S*-dependent segregation leading to the generation of a variable, even or odd number of offspring.

## Results

### *The chromosome of G1 predator cells features a longitudinal* ori – ter *arrangement with novel polarity*

To gain insight into the spatial organization of the chromosome in living *B. bacteriovorus* cells, we labeled the chromosomal origin (*ori*) and terminus (*ter*) using orthologous *parS*/ParB pairs (39). Here, the *parS*_*PMT1*_ or *parS*_*P1*_ sequence was integrated in the chromosome of the wild-type HD100 strain near to the predicted *ori* or *ter* locus, respectively (see Methods), and fluorescent fusions to the cognate *parS-* binding proteins ParB_PMT1_ and ParB_P1_ were constitutively produced from a replicative plasmid (Figure 2A). We first monitored the subcellular position of *ori* (YFP-ParB_PMT1_) or *ter* (CFP-ParB_P1_) in living attack-phase (AP) cells, using epifluorescence microscopy. Each locus was detected as a single unipolar focus in most cells (Figure 2B, Figure Supplement 1B), supporting the long-standing notion that AP cells carry only one copy of their chromosome and represent the non-proliferative (G1) phase of the cell cycle (5,27,38) (Figure 1). Similar analyses of fluorescence profiles in a series of control strains confirmed the specificity of each *parS*/ParB pair (Figure Supplement 1A, C), consistent with previous locus labeling data on other species (15,17,39–42). We found that *ori* and *ter* occupy opposite poles in the majority of the cells in which both loci were labeled (67% cells, n = 3593 in a representative experiment, Figure 2C, Figure Supplement 2C). To determine the polarity of this *ori-ter* arrangement, we took advantage of RomR, an essential component of a predation complex at the invasive pole of *B. bacteriovorus* (43) (Figure Supplement 2D). Interestingly, the *ter* marker localized at the pole opposite a RomR-tdTomato fusion in most cells (79% cells, n = 6061 in a representative experiment, Figure 2D). Consistently, images of G1 *B. bacteriovorus* cells attached to *E. coli* cells showed that *ori* occupies the invasive, non-flagellated pole, whereas *ter* localizes at the flagellated pole in the majority of cells (Figure 2E, Figure Supplement 2A). Additional evidence for this orientation was obtained using the sheathed unipolar flagellum (44) as a polarity beacon, labeled with membrane dyes in cells carrying the *ori* and/or *ter* labeling system (Figure 2F, Figure Supplement 2B-C, Supplementary Table 1). Taken together, these data show that in most G1 *B. bacteriovorus* cells, chromosomal *ori* and *ter* loci are positioned at the invasive and flagellated poles, respectively. Note that the preferential localization of *ter* is more flexible (at the flagellated pole in ~70% of cells regardless of the polar marker) compared to the strict localization of *ori* at the invasive pole. Strikingly, *B. bacteriovorus* features an inverse chromosomal polarity compared to other species carrying one (or more) unipolar flagellum, in which the chromosomal centromere is always located at the flagellated pole of newborn cells (e.g. *Caulobacter crescentus* (11,45)*, Vibrio cholerae* (46)*, Agrobacterium tumefaciens* (16)).

**Figure 2.**
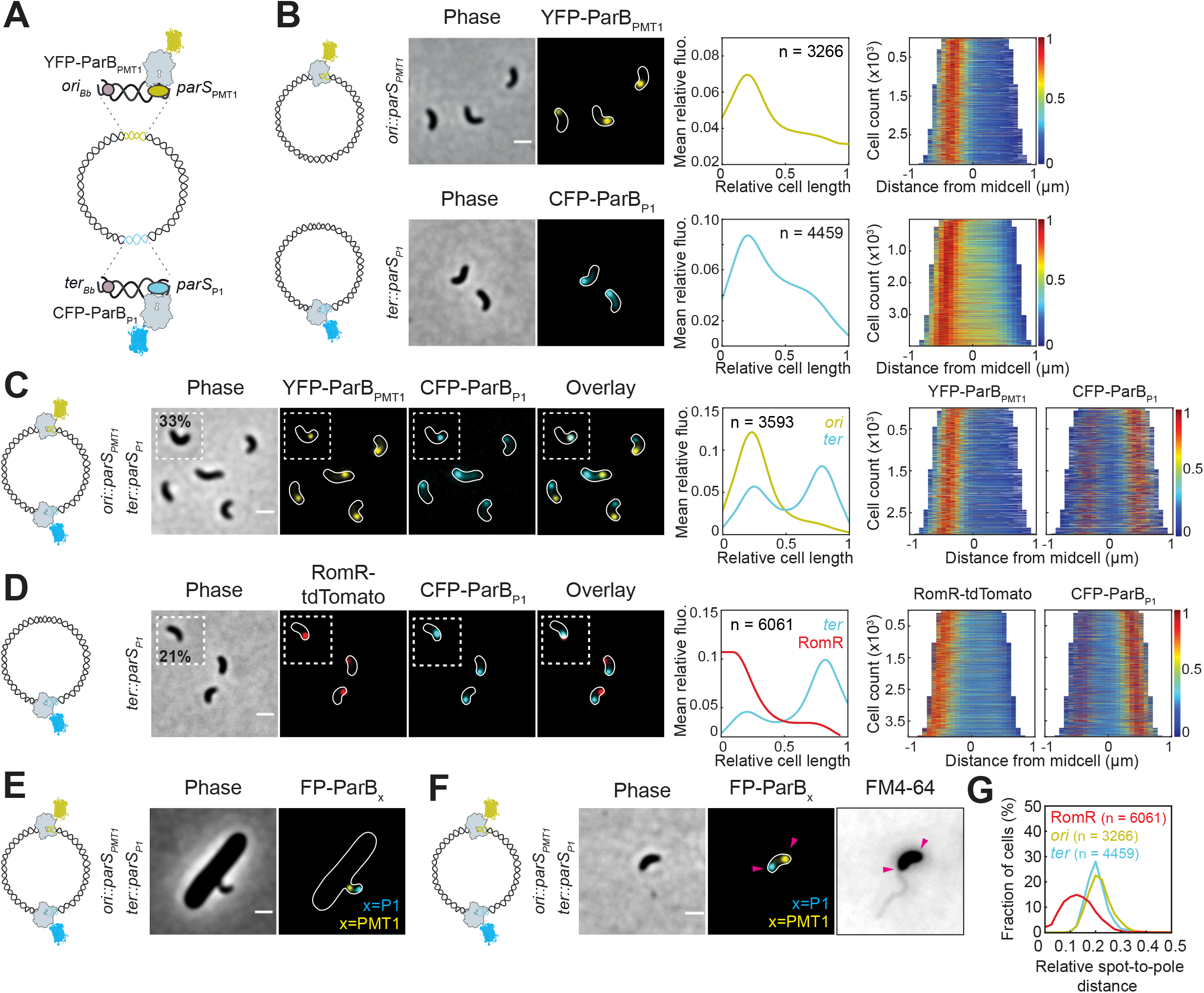
Linear arrangement of the *B. bacteriovorus* chromosome and localization of *oriC* near the invasive pole. **(A)** Schematics of the orthologous *parS-* ParB pairs used in this study to label chromosomal loci. The *parS*_*PMT1*_/*parS*_*P1*_ sequences (yellow/cyan) were integrated near the origin of replication (*ori*_*Bb*_) or terminus of replication (*ter*_*Bb*_) and the cognate YFP-ParB_PMT1_ or CFP-ParBs_P1_ fluorescent fusions were constitutively produced from a replicative plasmid, respectively. Strains carrying *parS*_*PMT1*_ or *parS*_*P1*_ are respectively referred to as *ori::parS*_*PMT1*_ or *ter::parS*_*P1*_. Loci where *parS*_*PMT1*_ or *parS*_*P1*_ were integrated are respectively referred to as *ori* or *ter.* **(B)** *ori* and *ter* are polarly localized. Left to right: representative phase contrast and fluorescence images of AP cells of *ori::parS*_*PMT1*_ and *ter::parS*_*P1*_ strains expressing cognate YFP-ParB_PMT1_ and CFP-ParBs_P1_ (GL868 and GL771, respectively); mean pole-to-pole profiles of relative fluorescence intensity of the corresponding fusion in the same cells; demographs of the corresponding fluorescent signal in the same cells sorted by length and oriented based on signal intensity. Heatmaps represent relative fluorescence intensities. **(C)** *ori* and *ter* occupy opposite poles in most cells. Left to right: representative phase contrast and fluorescence images of AP cells of a *ori::parS*_*PMT1*_ *ter::parS*_*P1*_ strain expressing cognate CFP-ParBs_P1_ and YFP-ParB_PMT1_ (GL995). Fraction of cells in which *ori* and *ter* colocalized is indicated (inset); fluorescence profiles as in B; demographs as in B, signals oriented based on YFP-ParB_PMT1_. **(D)** *ter* localizes at the pole opposite RomR in most cells. Left to right: representative phase contrast and fluorescence images of AP cells of *ter::parS*_*P1*_ strain expressing RomR-TdTomato and CFP-ParBs_P1_ (GL816). Fraction of cells in which RomR and ParBP1 colocalized is indicated (inset); fluorescence profiles as in B; demographs as in B, signal oriented based on RomR-TdTomato. **(E)** *ori* occupies the invasive pole and *ter* occupies the flagellated pole during prey attachment. Representative phase contrast and fluorescence images of AP cells of *ori::parS*_*PMT1*_ *ter::parS*_*P1*_ strain expressing cognate CFP-ParBs_P1_ and YFP-ParBPMT1 (GL995) 30 min after mixing with prey. **(F)** *ori* occupies the non-flagellated pole and *ter* occupies the flagellated pole. Representative phase contrast and fluorescence images of AP cells of *ori::parS*_*PMT1*_ *ter::parS*_*P1*_ strain expressing cognate CFP-ParBs_P1_ and YFP-ParB_PMT1_ (GL995) after staining with FM4-64. Arrowheads point at *ori* and *ter* foci at opposite poles. **(G)** Histogram of the relative distance from fluorescent spot of RomR-TdTomato (red), CFP-ParBs_P1_ (cyan) and YFP-ParB_PMT1_ (yellow) to the nearest cell pole for cells in B and D. For all panels, schematics illustrate the relevant *ori* and *ter* labelling construct. Scale bars are 1μm. n indicate the number of cells analysed in a representative experiment. For all, experiments were performed at least twice. For all panels, cell outlines were obtained with Oufti.

### *The chromosome of* B. bacteriovorus *is packed in a dense nucleoid that partially excludes freely-diffusing proteins*

Demographs and fluorescence profiles representing the subcellular localization of *ori* and *ter* loci (Figure 2B) or the polar marker RomR, (Figures 2D) suggest that *ori* and *ter* occupy slightly off-pole positions. Indeed, these loci were more distant from the closest cell pole than RomR (Figure 2G, Supplementary Table 2). This is consistent with the idea that the chromosome of *B. bacteriovorus* forms a nucleoid (*i.e.* not filling the entire cytoplasm) as previously proposed (47,48). Yet, this aspect of the *B. bacteriovorus* chromosome was never explored in living cells. Staining the DNA of G1 cells with fluorescent dyes confirmed the existence of a well-defined nucleoid occupying a fraction of the cytoplasm (Figure 3A-C, Figure Supplement 3A-C). The localization of *ori and ter* at the tips of the nucleoid (Figure 3D, Figure Supplement 3B) confirms the longitudinal arrangement of the *B. bacteriovorus* chromosome. Interestingly, nucleoid area was much smaller in G1 *B. bacteriovorus* cells than in *E. coli* (Figure 3A, E), despite roughly similar genome sizes (3.8 Mb and 4.7 Mb, respectively), showing that the chromosome of *B. bacteriovorus* is extremely compacted. Remarkably, we found that freely diffusing fluorescent proteins were at least partially excluded from the region of the cytoplasm occupied by the chromosome (Figure 3F), regardless of the fluorescent protein or the DNA dye used (Figure Supplement 3C, D). While it is well reported that large molecular complexes such as ribosomes (49–51), protein aggregates (52–54) or higher order assemblies (55,56) are excluded from nucleoids, this is the first time, to the best of our knowledge, that nucleoid exclusion is reported for relatively small, monomeric proteins. Our data thus reveal an outstanding degree of compaction of the *B. bacteriovorus* chromosome during the G1 stage of the cell cycle.

**Figure 3.**
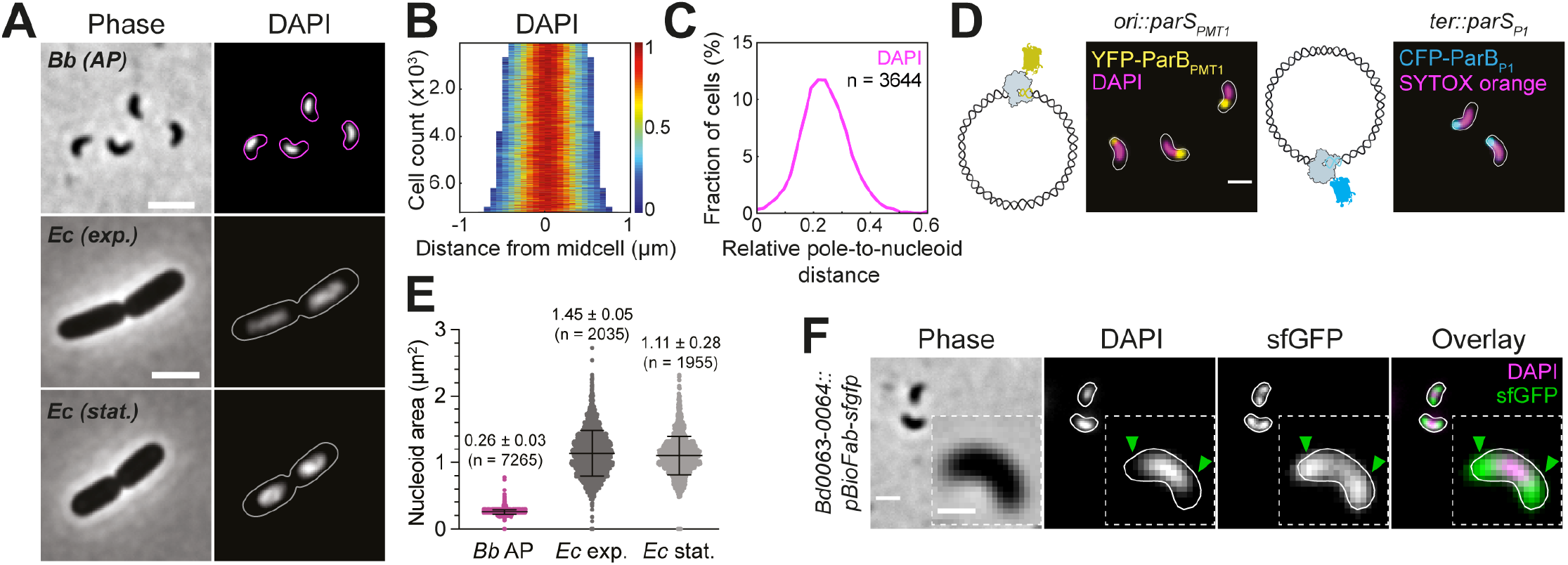
Nucleoid of *Bdellovibrio bacteriovorus* is organized in a dense nucleoid that partially excludes freely diffusing proteins. **(A)** The chromosome of *B. bacteriovorus* is organized in a well-defined nucleoid. From top to bottom: representative phase contrast and fluorescence images of fresh AP cells of WT *B. bacteriovorus* and in exponential and stationary phase cells of WT *E. coli* stained with DAPI (HD100 and MG1655, respectively). Scale bar is 2 μm. **(B)** Demograph of the DAPI signal from *B. bacteriovorus* images in A. Heatmap represent relative fluorescence intensities. **(C)** Relative pole-to-nucleoid distances for cells in B. n indicate the number of cells analysed in a representative experiment. **(D)** *ori* and *ter* colocalize with the nucleoid tips. Representative overlay images of AP cells of *ori::parS*_*PMT1*_ and *ter::terS*_*P1*_ strains expressing cognate YFP-ParB_PMT1_ or CFP-ParBs_P1_ (GL868 and GL771, respectively) stained with DAPI and SYTOX orange as indicated. Schematics illustrate the locus labelling construct. Scale bar is 1μm. **(E)** The nucleoid of *B. bacteriovorus* is extremely compacted. Distributions of nucleoid areas in WT *B. bacteriovorus* and exponential or stationary phase *E. coli*. Mean and standard deviation values are shown on top of the corresponding plot. n indicate the number of cells analysed in a representative experiment. **(F)** The nucleoid of *B. bacteriovorus* partially excludes freely diffusing proteins. Representative phase contrast and fluorescence images of AP cells constitutively expressing *sfgfp* from the *Bd0063-0064* intergenic locus (GL1212) stained with DAPI. Inset: enlarged example, arrowheads point to nucleoid exclusions. Scale bar is 1 μm except in the inset (0.5 μm). For all panels, cell outlines were obtained with Oufti.

### Spatial arrangement of the chromosome is maintained during the G1-S transition and DNA replication initiates at the invasive pole

We subsequently set out to investigate chromosome dynamics further in the cell cycle, when the predator cell resides within its prey, where it is expected to replicate its genomic content. To directly track DNA replication in living cells, we first monitored the subcellular distribution of DnaN, the replisome β-clamp commonly used as a proxy for replisome assembly and dynamics (57). We designed a scarless *dnaN*::*dnaN*-*msfgfp* construct at the native chromosomal locus in a wild-type background, allowing the production of a DnaN-msfGFP fusion in place of the endogenous DnaN (Figure Supplement 10). DnaN-msfGFP signal was diffuse in the cytoplasm in G1 cells (exhibiting the partial nucleoid exclusion described above) (Figure 4A), indicative of unassembled replisome and consistent with the absence of DNA replication in G1. A DnaN-msfGFP focus appeared at one cell pole 95 ± 14 minutes after mixing with an *E. coli* prey cell (from 3 independent time-lapse experiments, total *n* = 318 bdelloplasts; Figure 4B, Figure Supplement 8B), indicative of replisome assembly and DNA replication initiation. We define the period between prey entry and initiation of DNA replication, marked by a DnaN-msfGFP focus, as the G1-S transition. During that period, the chromosome of *B. bacteriovorus* was still compact and *ori*-*ter* polarity was maintained (Figure 4C). Importantly, we obtained several lines of evidence providing unambiguous support to the previously proposed idea that DNA replication initiates at the invasive pole (38): (i) fluorescent foci of DnaN and RomR fusions occupied the same pole (98% cells, n = 133; Figure 4D), (ii) the DnaN-msfGFP focus colocalized with *ori* (Figure 4D), which was labeled either with *parS*_*PMT1*_•YFP-ParB_PMT1_ (colocalization in 99% cells, n = 104) or with the centromeric protein ParB_Bb_ (see below; colocalization in 93% cells, n = 111), and (iii) Click™-labeling of the thymidine analog EdU revealed a DNA synthesis spot that colocalized with the *ori* region (Figure 4E, Figure Supplement 4A-C). These results also validate the use of the *parS*_*PMT1*_•ParB_PMT1_ pair as proxy for the chromosomal *ori* in *B. bacteriovorus.* Thus, the spatial arrangement of the chromosome seen in G1 cells is preserved during the G1-S transition upon prey invasion.

**Figure 4.**
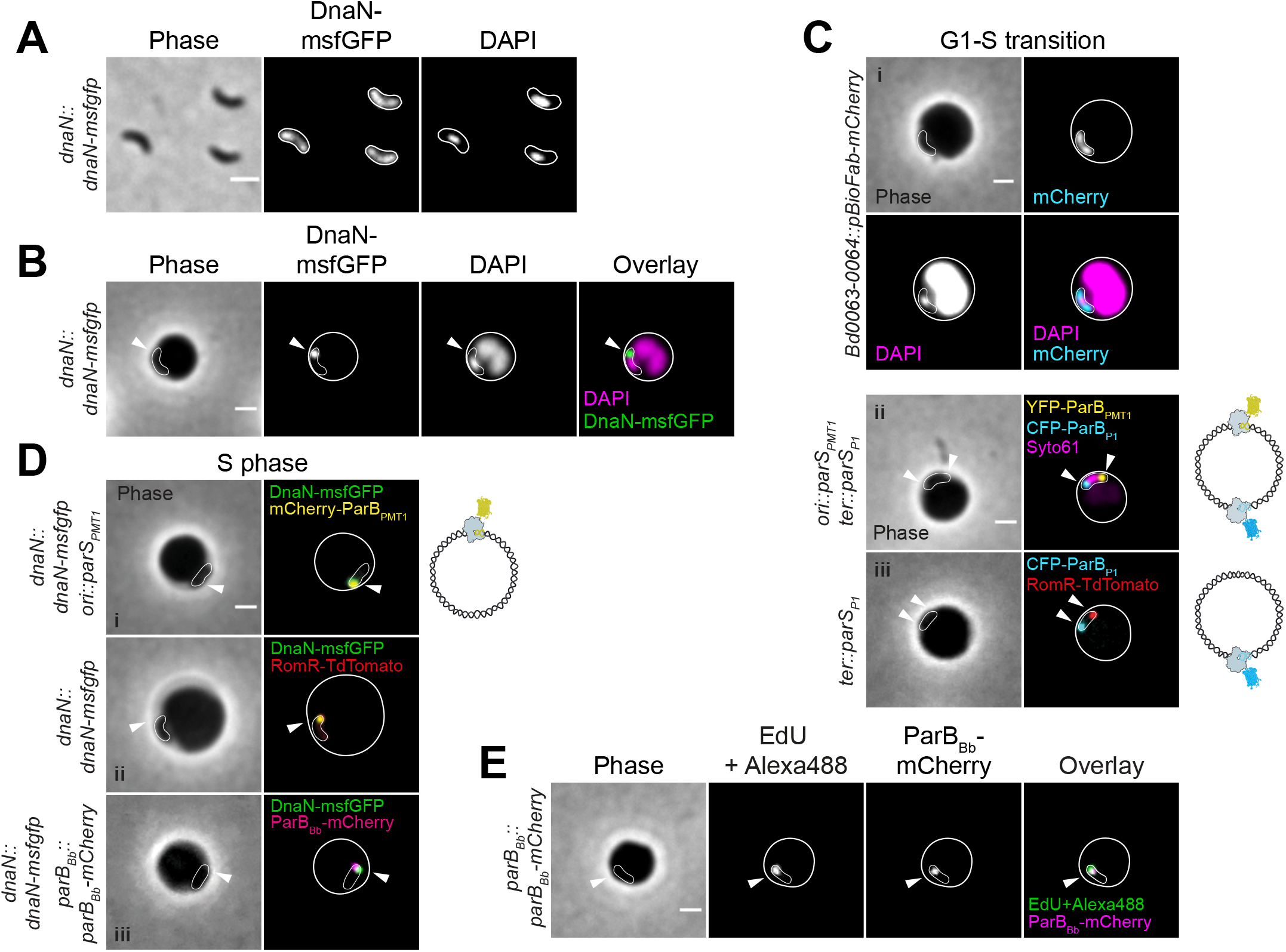
Chromosomal arrangement is maintained during the G1-S transition and DNA replication starts at the invasive pole. **(A)** DNA replication is inhibited in G1 phase and the diffuse DnaN signal exhibits partial nucleoid exclusion. Representative phase contrast and fluorescence images of AP cells of a *dnaN::dnaN-msfgfp* strain (GL673); cell outlines were obtained with Oufti. **(B)** Replisome assembles at one cell pole. Representative phase contrast and fluorescence images of *dnaN::dnaN-msfgfp* strain (GL673) stained with DAPI 60 min after mixing with *E. coli* prey. Arrowhead points to the DnaN-msfGFP focus indicative of an active replisome. **(C)** The chromosome of *B. bacteriovorus* is still compacted during G1-S transition and *ori-ter* polarity is maintained. Representative phase contrast and fluorescence images of (i) cells constitutively producing mCherry from a chromosomal locus (GL1025) stained with DAPI and imaged 30 min after mixing with *E. coli* prey, showing that the nucleoid does not expand to the cell poles unlike free mCherry used to visualize the whole cell; (ii) *ori::parS*_*PMT1*_ *ter::parS*_*P1*_ strain expressing cognate CFP-ParBP1 and YFP-ParB_PMT1_ (GL995) stained with Syto61 and imaged 35 min after mixing with *E. coli* prey; *ori* and *ter* occupied opposite poles (arrowheads) in 71% of cells (n=96 from one representative experiment); (iii) *ter::parS*_*P1*_ strain expressing cognate CFP-ParBs_P1_ and RomR-TdTomato (GL816) imaged 35 min after mixing with *E. coli* prey; *ter* and RomR occupied opposite poles in 73% of cells (n=77 from one representative experiment). **(D)** DNA replication initiates at the invasive cell pole (start of the S phase). Representative phase contrast and fluorescence images of (i) *ori::parSPMT1 dnaN::dnaN-msfgfp* strain expressing cognate mCherry-ParBPMT1 (GL1003) imaged 75 min after mixing with *E. coli* prey; colocalization in 99% of cells (n=104 from one representative experiment); (ii) *dnaN::dnaN-msfgfp* strain expressing RomR-TdTomato (GL1211) imaged 60 min after mixing with *E. coli* prey; colocalization in 98% of cells (n=133 from one representative experiment); (iii) *dnaN::dnaN-msfgfp parBBb::parBBb-mCherry* strain (GL1055) imaged 110 min after mixing with *E. coli* prey; colocalization in 93% (n=111 from one representative experiment). Colocalization was quantified manually. **(E)** Colocalization of newly synthesized DNA and the *oriC* region marked with the endogenous ParBBb at the beginning of the S phase in *B. bacteriovorus.* Representative phase contrast and fluorescence images of cells of *parBBb::parBBb-mCherry* strain (GL906) 150 min after mixing with prey, exposed to a 5 minute pulse of the nucleotide analogue EdU, which was fluorescently labelled with Alexa488. ParBBb-mCherry was used to label *oriC* since DnaN-msfGFP foci were unstable in this experimental setup. Arrowheads point to fluorescent foci. Schematics illustrate the *ori* and *ter* labelling construct used in each panel. *B. bacteriovorus* and bdelloplasts outlines in panels B-E were drawn manually based on the phase contrast images. Scale bars are 1 μm.

### Replisome dynamics reveal multiple concomitant rounds of DNA replication in the growing filament

Examination of hundreds of predator cells imaged in time-lapse upon DNA replication initiation revealed common subcellular patterns. First, the DnaN-msfGFP focus migrated from the invasive pole to a midcell position (Figure 5A, arrowhead), likely reflecting the progression of replication along the mother chromosome (58,59). A second DnaN-msfGFP focus was detected 167 ± 15 minutes post-mixing with prey, *i.e.* 71 ± 5 min after the first focus, usually at the invasive pole (in 73,9% of 318 bdelloplasts from 3 independent time-lapse experiments) (Figure 5A, arrow, Figure Supplement 8B). Transient splitting or merging of foci was occasionally observed, possibly representing the two replication forks, as reported in *Caulobacter crescentus* (60). A third DnaN-msfGFP spot formed at the opposite pole (Figure 5A, asterisk), followed by additional DnaN-msfGFP foci, which showed highly dynamic movements, indicating that more than two replisomes are simultaneously active in the growing predator cell (Figure 5A, Movie 1). Similar dynamics were obtained when a fluorescent fusion of the clamp-loader component DnaX was used to label the replisome, as done in other species (61–64), although the signal intensity of DnaX-msfGFP (replacing the native DnaX, *dnaX::dnaX-msfgfp*) was weaker than the DnaN fusion (Figure Supplement 5A). The gyrase inhibitor novobiocin, which specifically blocks DNA replication initiation (65), prevented the formation of the first or subsequent DnaN-msfGFP foci depending on when the drug was added in the time course of the cell cycle (Figure Supplement 5B). Thus, our data show that the number of replisomes increase in the growing *B. bacteriovorus* and that multiple rounds of DNA replication can occur concomitantly.

**Figure 5.**
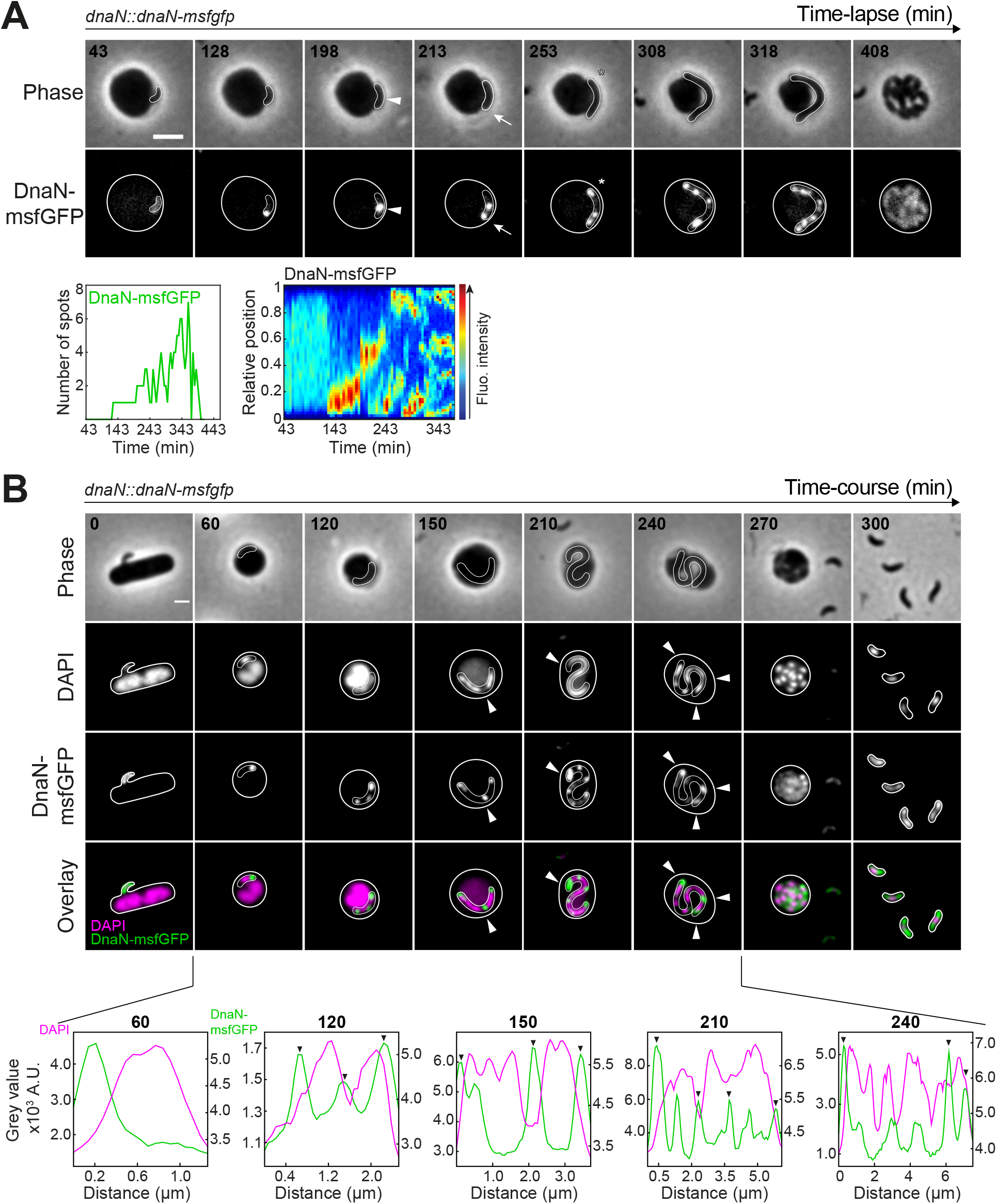
Multiple concomitant rounds of DNA replication in *B. bacteriovorus.* **(A)** Replisome dynamics during the proliferative (S) phase. *B. bacteriovorus* strain GL673 (*dnaN::dnaN-msfgfp*) was mixed with prey and imaged in time-lapse after 43 min at 5 min intervals. Top: phase contrast and fluorescence images of selected timepoints of a representative experiment are shown. Arrowhead points to a mid-cell positioned DnaN-msfGFP focus; arrow points to the second DnaN-msfGFP focus; asterisk points to the third DnaN-msfGFP focus. The full time-lapse is shown in Movie 1. Bottom: number of DnaN-msfGFP spots detected in Oufti over time (left) and kymograph of the DnaN-msfGFP signal along the cell length (right), for the same cell. Scale bar is 2 μm. **(B)** Nucleoid decondenses after DNA replication initiation. *B. bacteriovorus* strain GL673 (*dnaN::dnaN-msfgfp*) was mixed with prey and imaged in time-course at 30 min intervals. Top: phase contrast and fluorescence images of selected timepoints from a representative experiment are shown; arrowheads point to regions with less DAPI signal and where replisomes are located. Scale bar is 1 μm. For all, outlines of *B. bacteriovorus* and bdelloplasts were drawn manually based on phase contrast images. Bottom: fluorescence intensity profiles of the corresponding signals in the same cells; arrowheads point to regions with less DAPI signal and where replisomes are located.

### Nucleoid decompaction occurs after DNA replication initiation

The extreme nucleoid compaction in G1 cells questions the accessibility of the chromosome to the machinery in charge of its replication. Highly dynamic replication factories (Figure 5A) suggest that the chromosomal copies are not as densely compacted in the S phase compared to the G1 and G1-S phases. To gain insight into the predator’s nucleoid compaction during growth inside the prey, we imaged DAPI-stained cells of a strain constitutively producing free mCherry (to label the whole predator cell), during a time-course of prey infection. The nucleoid occupied a restricted area in the cell (compared to the mCherry signal) even when the first DnaN-msfGFP focus was detected (Figure 4B, Figure Supplement 5C). At later stages when additional replisomes were active, the DAPI signal filled larger cell areas, indicating at least partial nucleoid expansion (Figure 5B, timepoint 120 min, Figure Supplement 5C, timepoint 120 min). Strikingly, areas that were the least stained with DAPI, presumably corresponding to regions of lower DNA density, were often occupied by a DnaN-msfGFP-labeled replisome (Figure 5B, arrowheads). Thus, nucleoid decompaction occurs, at least partially, after DNA replication initiation. Nucleoid segregation (marked by distinct DAPI-stained units) was visible at late time points (Figure 5B, arrowheads, Figure Supplement 5C, asterisks). Thus, we propose that nucleoid de-condensation, possibly triggered by the progression of the replisomes, is followed by re-compaction of the chromosomes, which happens before cell division.

### *Duplicated* ori *undergo asymmetric polar segregation and* ter *segregation is uncoupled from cell division*

The simultaneous occurrence of multiple replication events in the growing predator cell raises the question of how and when the newly synthesized chromosomes are segregated. Since the subcellular positions of *ori* and *ter* loci determine the dynamics of chromosome segregation in other species (7), we examined the spatial arrangement of the chromosome during the proliferative (S) phase of the cell cycle. Time-lapse imaging of labeled *ori* showed that a second focus appeared and quickly moved towards the opposite pole (Figure 6A, Figure Supplement 6A, Movie 2), reminiscent of the asymmetric *ori* segregation described in several binary-dividing species (11), Deghelt:2014ej, Dubarry:2019fu, Fogel:2006km, Frage:2016eq}. In line with this model (23,45), the *ter* focus shifted from its polar position to midcell (Figure 6B, E, Figure Supplement 6B). Interestingly, this first round of segregation was achieved to completion, as the *ter* copies clearly split in two distinct foci at midcell (Figure 6B, E, Movie 3). Thus, *ter* segregation is temporally uncoupled from cell constriction. To further investigate segregation dynamics, we turned to the endogenous ParB (here named ParB_*Bb*_) to mark the native centromeric *parS_Bb_* site located next to *ori* (Figure Supplement 1A). Consistent with the reported biphasic expression pattern of the corresponding operon (66,67), we could only detect weak fluorescent signal in G1 cells of a strain in which *parB_Bb_* was replaced by *parB_Bb_-mcherry* (Figure Supplement 6C). Nevertheless, while no detectable foci could be observed in the G1 phase (see below), ParB_*Bb*_-mCherry always formed a focus that colocalized with the first DnaN-msfGFP-labeled replisome (Figure 4D). Furthermore, the first duplication and polar segregation of ParB_*Bb*_-mCherry foci showed similar dynamics as *parS_PMT1_•*YFP-ParB_PMT1_-labeled *ori* (Figure 6A, Figure 6C, Figure Supplement 6D), supporting our previous finding of asymmetric *ori* segregation and the use of ParB_*Bb*_-mCherry as *ori* marker. Monitoring the dynamics of both ParB_*Bb*_ and *ter* in the same cells showed that *ter* relocation to midcell started after the newly duplicated *ori* reached the non-invasive pole (Figure 6E, time point 200 min).

**Figure 6.**
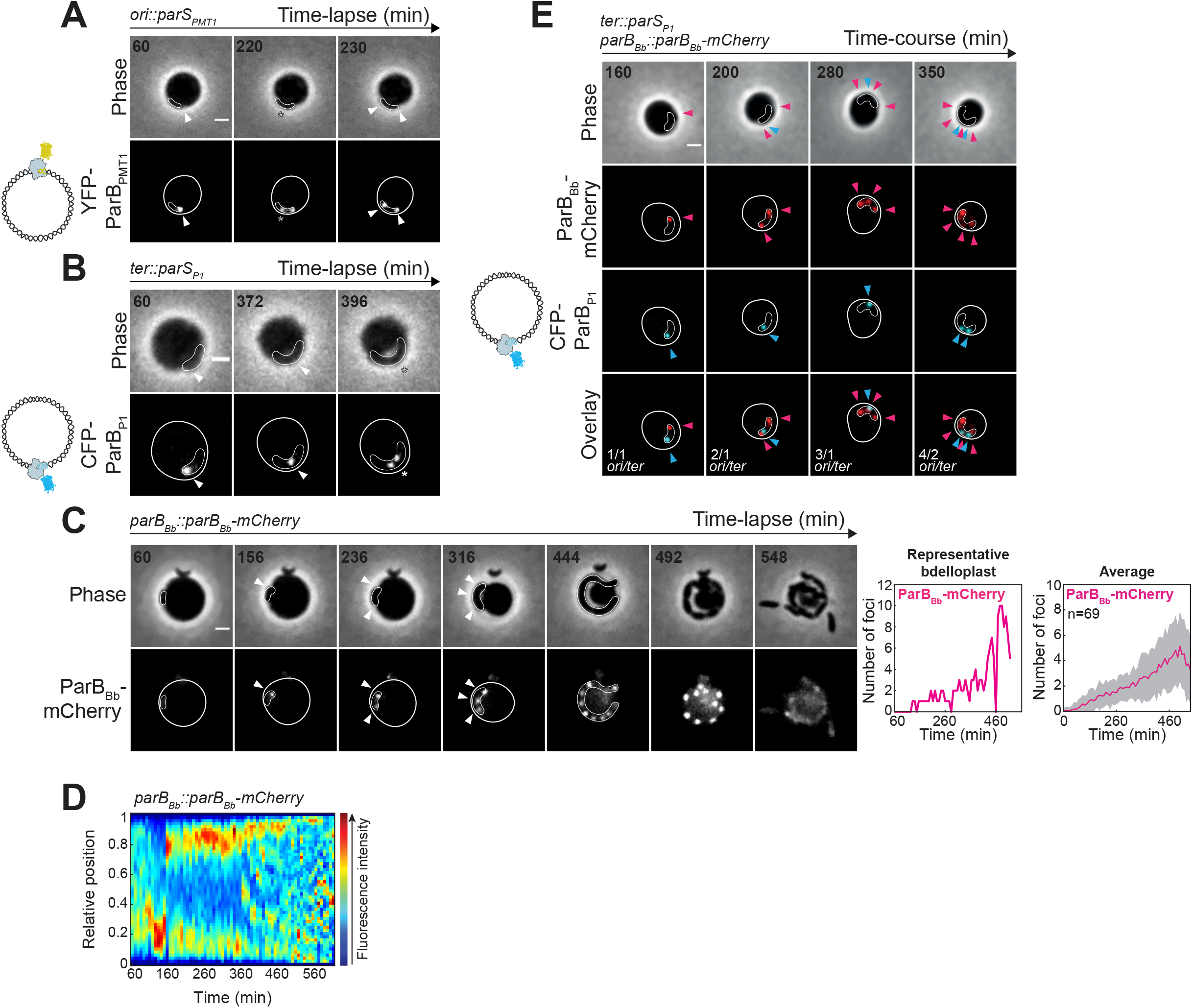
Spatio-temporal arrangement of the chromosome during the proliferative phase of the cell cycle in *B. bacteriovorus.* **(A)** First round of *ori* segregation is asymmetric. *B. bacteriovorus* strain *ori::parS*_*PMT1*_ expressing cognate YFP-ParB_PMT1_ (GL868) was mixed with prey and imaged in time-lapse after 60 min with 10 min intervals. Phase contrast and fluorescence images of selected timepoints are shown; arrowheads point to the *ori* focus before replication and the duplicated *ori* foci after segregation; asterisk points to the *ori* copy being segregated towards the opposite pole. The time-lapse is shown in Movie 2. **(B)** *ter* segregation is temporally uncoupled from cell division. *B. bacteriovorus* strain *ter::parS*_*P1*_ expressing cognate CFP-ParBs_P1_ (GL771) was mixed with prey and imaged in time-lapse after 60 min with 8 min intervals. Phase contrast and fluorescence images of selected timepoints are shown; arrowhead and asterisk point to mid-cell *ter* localization and segregation, respectively. The time-lapse is shown in Movie 3. **(C)** ParB-driven *ori* segregation during S phase. *B. bacteriovorus* strain GL906 (*parBBb:: parBBb-mCherry*) was mixed with prey and imaged in time-lapse after 60 min with 8 min intervals. Left: phase contrast and fluorescence images of selected timepoints are shown. Arrowheads point to the first ParBBb-mCherry focus, two segregated ParB foci, then 3 segregated foci; multiple well-separated ParBBb-mCherry foci are visible at time-point 444 min. The time-lapse is shown in Movie 4. Right: number of ParBBb-mCherry spots detected in Oufti, over time for the same representative bdelloplast; average number of ParBBb-mCherry spots detected in Oufti, over time; grey area indicates standard deviation; n indicates the average number of bdelloplasts analysed until timepoint 384 min, from which the number of bdelloplasts that could be analysed progressively decreased to 62. **(D)** Kymograph of the ParBBb-mCherry signal along the cell length for one representative cell. Arrowheads indicate timing of, from left to right, the first ParBBb-mCherry focus, pole-to-pole segregation upon duplication, and the third focus. **(E)** *ter* relocation to mid-cell starts after the second *ori* reaches the non-invasive pole. *B. bacteriovorus* strain *parBBb::parBBb-mCherry ter::parSP1* expressing cognate CFP-ParBP1 (GL1368) was mixed with prey and imaged in time-course with 30 min intervals. Phase contrast and fluorescence images of selected timepoints are shown; pink arrowheads point to *ori* copies; blue arrowheads point to *ter* copies. Schematics illustrate the *ori* and *ter* labelling constructs used in each panel. Scale bars are 1 μm. For all, outlines of *B. bacteriovorus* and bdelloplasts were drawn manually based on phase contrast images.

### Chromosome segregation occurs progressively as new copies are being synthesized

After the first “binary-like” replication and segregation round, additional ParB_*Bb*_ foci gradually appeared (Figure 6C-D Figure Supplement 6D, Movie 4). The highest number of ParB_*Bb*_ foci varied between bdelloplasts and reached 5 on average under these conditions (*n* = 69, representative time-lapse experiment) (Figure 6C). As expected, the ParB_*Bb*_-mCherry foci always colocalized with YFP-ParB_PMT1_-labeled *ori* on snapshots taken at various times during the proliferative phase of the cell cycle (Figure Supplement 6E). Strikingly, ParB_*Bb*_ foci were always evenly distributed during filamentous growth (Figure 6C, Figure Supplement 6D, Movie 5), indicating that even after the first round of replication, *ori* segregation occurs as soon as new chromosomal copies are being synthesized. In addition, temporal uncoupling of *ter* segregation from cell division was not limited to the first replication round, as additional *ter* foci appeared before visible cell constriction (Figure Supplement 6B, asterisks). We did not observe *ter* foci at the poles of the growing cell, unlike *ori* foci (Figure Supplement 6B, asterisks, Figure 6C, E), hinting that *ori* but not *ter* will occupy the two “old” poles transmitted from the mother cell to the progeny. In the last minutes before non-binary cell division, *ter*-associated signal dispersed, and foci reappeared in the progeny (Figure Supplement 6B, Movie 6), which could suggest a temporary reorganization of the *ter* macrodomain during the division process.

### *The ParABS system is required for progressive* ori *segregation and accurate cell cycle progression*

The asymmetric pole-to-pole segregation of the first duplicated centromere and the progressive partitioning of additional copies strongly suggest that the ParAB*S* system drives these segregation events in *B. bacteriovorus*. To examine this idea further, we introduced perturbations in that system by constitutively producing ParB_*Bb*_ fusions from a plasmid (Figure Supplement 10), which is expected to modify the ParA:ParB interplay (68,69). Overproduced ParB_*Bb*_-FP fusions formed distinct foci in predator cells after prey invasion (Figure 7A, arrowhead), which perfectly colocalized with the YFP-ParB_PMT1_-labeled *ori* copies (Figure Supplement 7D). However, the localization pattern of these ParB_*Bb*_•*ori* complexes (Figure 7A, asterisks, Movie 7) differed from cells in which ParB_Bb_ is produced at native levels (Figures 2, 6), consistent with segregation defects: (i) in longer cells the second ParB_*Bb*_ focus rarely reached the opposite pole and instead stalled in the middle of the cell (Figure 7A-B) and (ii) the number of foci did not regularly increase before division, and fusions of existing foci were observed (Figure 7A, time point 290 min). Moreover, constitutive production of ParB_*Bb*_, either untagged or in fusion with mCherry or msfGFP, led to pronounced phenotypes in the released progeny: (i) they displayed more variable and on average larger cell length and nucleoid area than control strains (Figure 7C; Figure Supplement 7B, Supplementary Table 3); (ii) cells had two or more *ori* foci (Figure Supplement 7C). Altogether, our data are consistent with the idea that the ParAB*S* system is required to achieve multiple progressive rounds of asymmetric *ori* segregation in *B. bacteriovorus*. Of note, nucleoid exclusion of cytoplasmic proteins was still observed in ParB_*Bb*_-overexpressing cells, indicating that the mechanism of chromosome compaction is independent of ParAB*S*-mediated chromosome segregation (Figure 7D).

**Figure 7.**
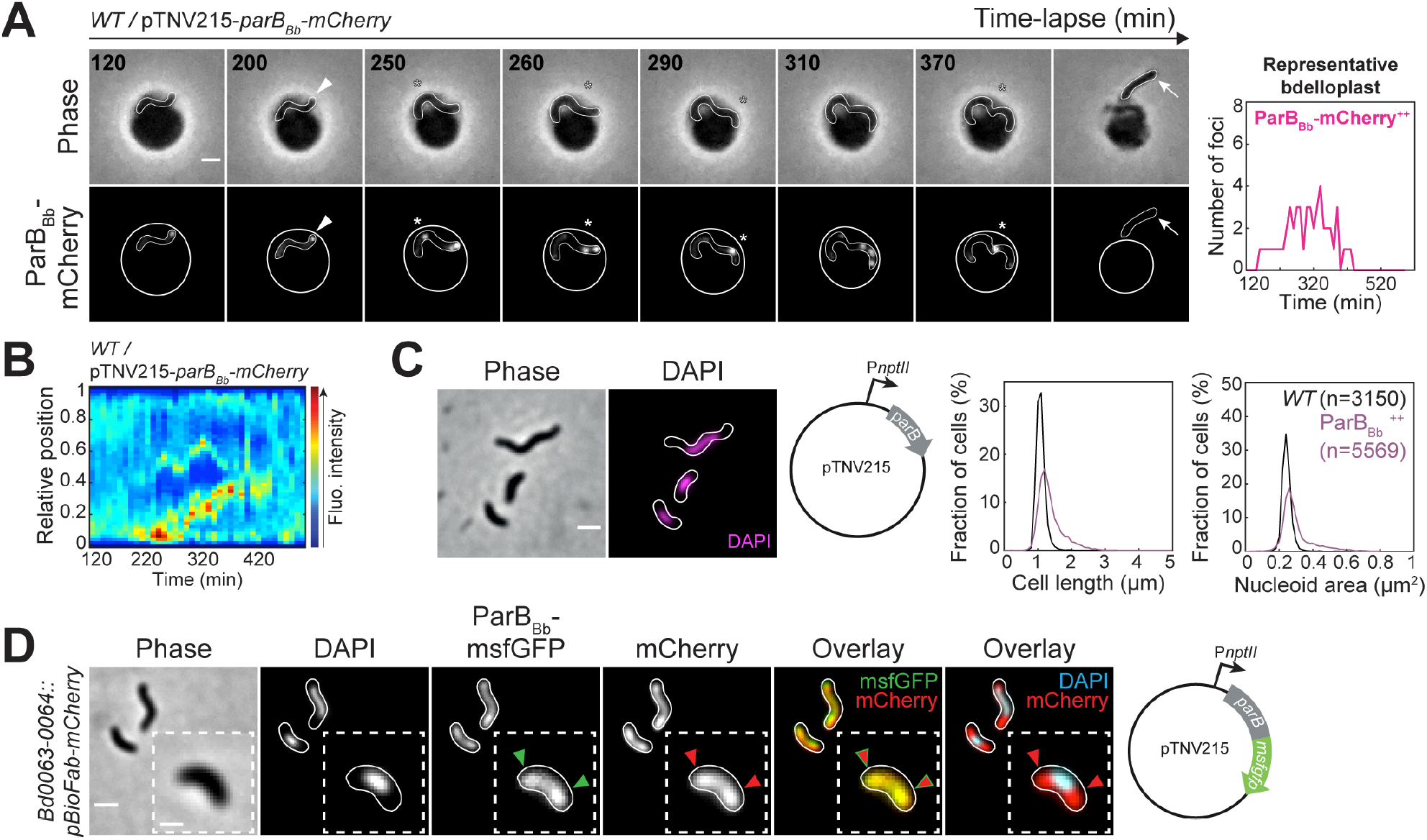
The ParABS system contributes to progressive *ori* segregation. **(A)** Overproduction of ParBBb leads to chromosome segregation defects. *B. bacteriovorus* strain GL1002 (*WT/*pTNV215-*parBBb-mCherry*) was mixed with prey and imaged in time-lapse after 120 min with 10 min intervals. Left: phase contrast and fluorescence images of selected timepoints; arrowhead points to a ParBBb-mCherry focus; asterisks point to altered *ori* behavior during the cycle; arrow points to a long daughter cell. The time-lapse is shown in Movie 7. Right: number of ParBBb-mCherry spots detected in Oufti, over time for the same representative bdelloplast. **(B)** Kymograph of the ParBBb-mCherry signal along the cell length for the same representative bdelloplast as in A. **(C)** Overproduction of ParBBb leads to phenotypic changes in AP cells. From left to right: representative phase contrast and fluorescence images of AP cells of a *WT* strain constitutively expressing untagged ParBBb (GL1261) stained with DAPI; histograms of cell length and nucleoid area for the same strain (purple) compared to *WT* (black); mean and standard deviation values are shown in Table 4. n indicates the number of cells analysed for each strain in a representative experiment. **(D)** Partial nucleoid exclusion of cytoplasmic fluorescent proteins in *parB*^++^ strain. Representative phase contrast and fluorescence images of cells constitutively producing mCherry from the *Bd0063-0064* intergenic locus and ParBBb-msfGFP from a plasmid (GL1388) stained with DAPI. Arrowheads point to nucleoid exclusions on an enlarged example (inset). Scale bars are 1 μm except for enlarged examples where scale bar is 0.5 μm. Outlines of *B. bacteriovorus* and bdelloplasts were drawn manually based on phase contrast images except in C where they obtained with Oufti. Experiments were performed at least twice.

### ParB_Bb_ binds the chromosomal centromere only after replication initiation

During these experiments, we surprisingly noticed that overproduced fluorescent ParB_*Bb*_ fusions displayed strong cytoplasmic signal but no focus in G1 cells (Figure Supplement 7A), suggesting that the above-mentioned absence of natively produced ParB*Bb-*mCherry focus (Figure Supplement 6C) could not be solely explained by low protein amounts. This finding raises the intriguing hypothesis that endogenous ParB_*Bb*_ is unable to accumulate at *parS_Bb_* sites during the G1 phase of the cell cycle. To get spatiotemporal insight into ParB_*Bb*_ focus formation relative to cell cycle progression, we constructed a strain producing both DnaN-msfGFP and ParB*Bb-*mCherry as single copies from their native chromosomal locus. Strikingly, ParB_*Bb*_ does not localize at *parS_Bb_* sites before the onset of DNA replication in *B. bacteriovorus,* since the ParB_*Bb*_-mCherry signal formed a first detectable focus after DnaN-msfGFP (41 minutes on average, from single-cell analysis in double-labeled strains, Figure 8A, Figure Supplement 8A; or when comparing population averages of single-labeled strains, Figure Supplement 8B-C). However, ParB_*Bb*_ does not require an active replisome to sustain accumulation at *parS_Bb_*, since we could still detect ParB_*Bb*_-mCherry foci at the end of the S phase when the DnaN-msfGFP foci disassembled (Figure 8A, Figure Supplement 8A, Movie 8). The ParB_*Bb*_-mCherry signal became diffuse after cell constriction started (Figure 6C, Figure 8A, Figure Supplement 8A), consistent with the absence of focus in G1 cells (Figure Supplement 6C). Thus, our data show that in *B. bacteriovorus,* ParB_*Bb*_ does not accumulate on its cognate *parS_Bb_* sites during the G1 phase and G1/S transition.

**Figure 8.**
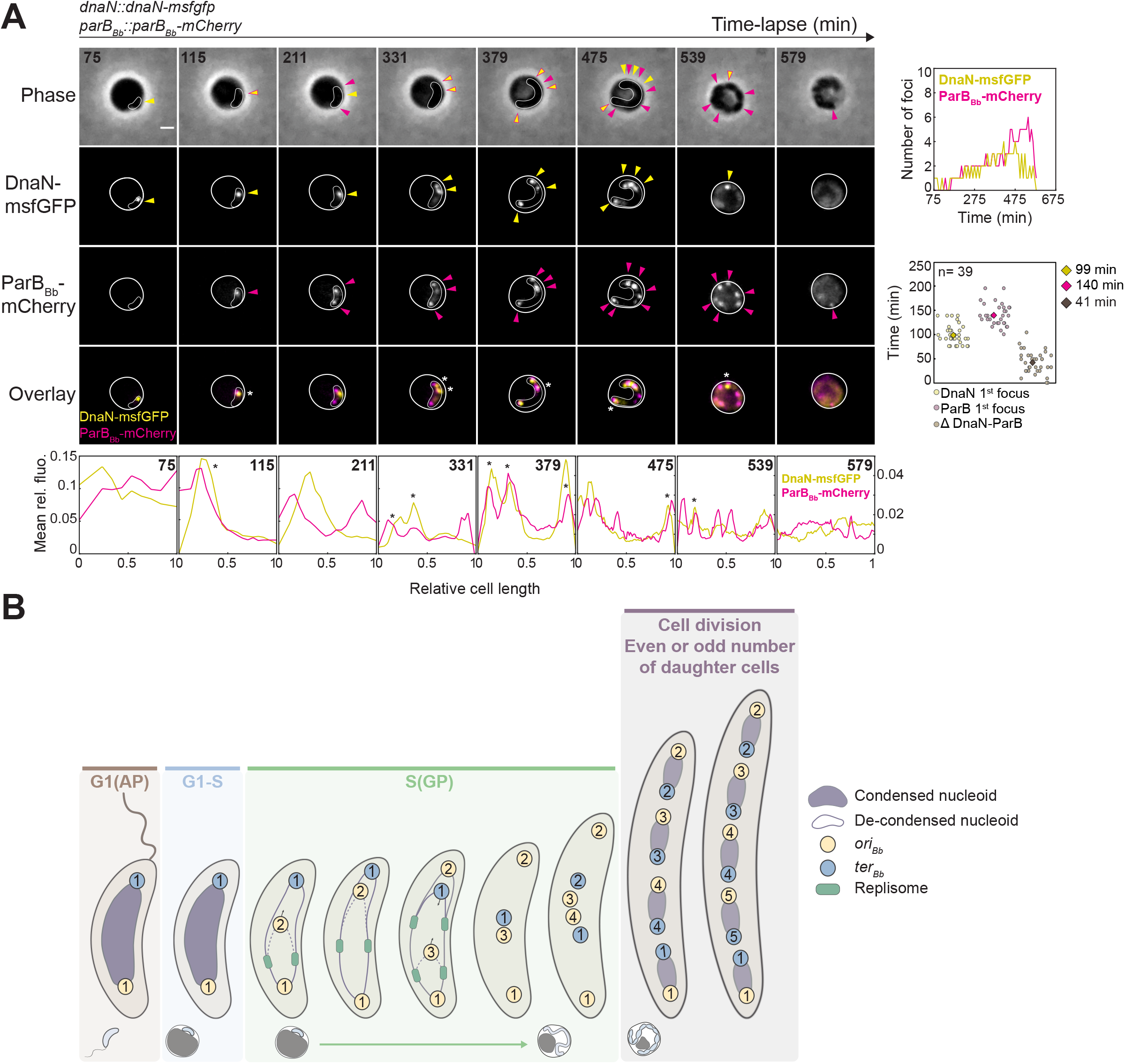
Stochastic chromosome replication initiation at multiple *oriC* and progressive segregation producing odd or even offspring. **(A)** ParBBb forms foci after DNA replication initiation and multiple copies of *oriC* can serve as a template for the next replication round. *B. bacteriovorus* strain GL1055 (*dnaN::dnaN-msfgfp parBBb::parBBb-mCherry*) was mixed with prey and imaged in time-lapse after 75 min with 8 min intervals. Top: left: phase contrast and fluorescence images of selected timepoints are shown. Arrowheads point to fluorescent foci; asterisks and two-colored arrowheads point to colocalization of the corresponding fluorescence signals. The time-lapse is shown in Movie 8; right: number of DnaN-msfGFP (yellow) and ParBBb-mCherry (pink) spots detected in Oufti, over time for the same representative bdelloplast; SuperPlot representation of the time of appearance of first DnaN-msfGFP and ParBBb-mCherry foci and the time difference between them, in single cells of the GL1055 strain; the average for each signal is represented as a colored diamond; n indicate the number of cells analysed in this experiment. Bottom: mean pole-to-pole profiles of relative fluorescence intensity of the corresponding fusions in the same cells; asterisks point to colocalization of the corresponding fluorescence signals. Scale bar is 1 μm. Outlines of *B. bacteriovorus* and bdelloplasts were drawn manually for display based on phase contrast images. **(B)** A model for non-binary chromosome choreography in *B. bacteriovorus.* The highly condensed nucleoid of G1 (AP) cells is longitudinally arranged such that *ori* (yellow) occupies the invasive pole, and more flexible *ter* (blue) occupies the flagellated pole. Once inside the prey, cells experience a G1/S transition during which the state of the chromosome is apparently unchanged. At the beginning of the proliferative S phase, DNA replication starts from the invasive pole and the duplicated *ori* is segregated asymmetrically. Nucleoid visibly decondenses after DNA replication initiation. When the 2^nd^ *ori* reaches the opposite pole, the replisome is at mid-cell. The 1^st^ *ter* then quickly moves from pole to mid-cell where it colocalizes with the 3^rd^ *ori* copy, which was newly synthesized and segregated (usually from the same invasive pole). Progressive *ori* and *ter* segregations continue, following new DNA replication rounds where variable numbers and copies of *ori* serve as initiation platform, leading to odd or even ploidy. Distinct nucleoids are visible again before division of the mother cell by multiple fission. Nucleoid schematic for last two cells in S phase is omitted for clarity.

### *Multiple* ori *copies serve as platforms for asynchronous replication rounds*

Finally, we asked how multiple chromosome replication events were orchestrated over time in growing *B. bacteriovorus* cells, *i.e.* whether DNA replication initiation steps were biased towards a specific subset of *ori* copies. Observation of bdelloplasts imaged in time-lapse showed that a DnaN-msfGFP spot colocalized with a ParB_*Bb*_-mCherry-labeled *ori* at several places in the cell (n = 65 in a representative experiment; Figure 8A, Figure Supplement 8A, asterisks), suggesting that (i) all concomitant replication rounds likely initiated from chromosomal *ori* loci, and do not represent ectopic replication events occurring from non-*ori* loci as reported for R-loops (70), and (ii) distinct *ori* loci can serve as replication initiation platforms. Although a fraction of DnaN-msfGFP foci often clustered near the cell ends (Figure 5A, Figure 8A, see Discussion), they were also observed in other cell regions. Consistently, the *ori* copies marked by both ParB_Bb_-mCherry and DnaN-msfGFP (probably representing *ori* being replicated) occupy diverse subcellular positions and vary in number over time and among bdelloplasts (Figure 8A, Figure Supplement 8A). Thus, we propose that the DNA replication initiation steps are asynchronous, or at least do not follow a readily predictable pattern beyond the first two rounds. This behavior results in the non-exponential increase in chromosome numbers, consistent with odd or even numbers of daughter cells being released at each generation (Figure 8B).

## Discussion

In this study, we benchmarked the use of key fluorescent reporters to monitor the subcellular dynamics of chromosomal loci as well as the replication and segregation machineries in living *B. bacteriovorus* cells. Semi-automated analysis of intracellular features at the single-cell and population levels allowed us to shed light on how the chromosome is organized in space and time during the cell cycle, opening the way for future quantitative cell biological approaches in this bacterium. Taken together, our data suggest a model (Figure 8B), in which asynchronous initiation of multiple DNA replication rounds is sufficient to elucidate why chromosome copies do not necessarily double at each replication round, leading to variable, odd or even numbers. Indeed, while several chromosomes simultaneously served as replication templates, usually not all of them were being copied at the same time. The observation that the invasive pole hosts the second round of replication in ~70% of cells suggests that the “mother” *ori* present at this pole is somehow primed, favoring re-initiation of chromosome replication at this location compared to the newly synthesized *ori*. Consistent with this idea, additional replisomes often accumulated from that pole in the growing predator, and later from the opposite pole, although DnaN foci were observed in other regions of the cell as well. What governs such spatial organization of DNA replication remains to be discovered. Considering the importance of transmitting a single and complete chromosome to each daughter cell, the dynamics of asynchronous replication initiation are likely not determined by chance. We think that complex regulation occurs in both space and time to prevent replication from all *ori’s* at the same time, and most importantly to avoid starting new synthesis that would prematurely end when prey resources are exhausted. Even though the temporal control of the S phase with respect to cell cycle progression and synchronous divisions is still unclear, our data provide insight into this question as we show that the late steps of chromosome segregation and cell division are uncoupled. In line with this idea, the positioning of one *ori* at both old poles of the pre-divisional mother cell inevitably results in at least one septum not being placed between two *ter* copies (Figure 8B).

Our study also revealed unique aspects of the spatial organization of *B. bacteriovorus* cells. First, the *ori* locus was always located near the invasive, non-flagellated pole of G1 *B. bacteriovorus* cells, thus in contrast with the chromosomal orientation architectures in previously characterized mono-flagellated bacteria (11,16,45,46). Based on RomR and flagellum labelling experiments, we conclude that the fraction of cells (~30%) in which *ori* and *ter* colocalized correspond to cases where *ter*, but not *ori*, is “misplaced”. Whether the occasional presence of *ter* at the invasive pole results from to the relative flexibility of the *ter* macrodomain (reported in *E. coli* (71,72)) or from another aspect related to non-binary proliferation remains to be discovered.

The dense *B. bacteriovorus* nucleoid unexpectedly alters the distribution of all reely-diffusing fluorescent proteins that we tested. So far, only larger objects were reported to be partially or fully excluded from the nucleoid (e.g. ribosomes or protein aggregates in *E. coli* (49–54)); these observations thus reveal an unprecedented degree of compaction of the *B. bacteriovorus* chromosome that constraints the diffusion of small molecules. Our data suggest at least partial de-condensation of the nucleoid during the S phase, matching with higher chromosome processing activities (including DNA replication but also transcription (66,73)), which may remodel the nucleoid and/or require increased accessibility within the nucleoid. Future work will aim at identifying the molecular and cellular factors that control the degree of nucleoid condensation, and its physiological impact with respect to the *B. bacteriovorus* cell cycle.

Finally, we found that ParB_*Bb*_ is unable to accumulate at the *ori* region during the G1 and G1-S transition stages, even when produced constitutively. The detection of ParB_*Bb*_ foci after DnaN foci suggests that DNA replication initiation might open up the *ori* region, allowing ParB_*Bb*_ binding. Besides, the accessibility of *parS_Bb_* may vary during the cell cycle depending on the level of nucleoid compaction; however, the heterologous *parS*_*PMT1*_ inserted near endogenous *parS_Bb_* sites remained accessible to ParB_PMT1_ at all times. To the best of our knowledge, this is the first example of a ParB homolog that does not permanently localize at the chromosomal centromere, regardless of protein levels, hinting that besides transcriptional modulation (66,67), ParB_Bb_ function may be subjected to novel cell cycle-dependent regulation.

Altogether, our data illustrate that *Bdellovibrio* is a treasure-trove for future discoveries of novel cell cycle regulation and cellular organization strategies. Moreover, our study sets the path for using *B. bacteriovorus* as a model to expand the quantitative investigation of subcellular events in bacteria, and highlights the exploitation of conserved proteins to address the needs of complex non-binary reproduction.

## Material and methods

### Strains

All strains and plasmids used in this study are listed in Supplementary Table 4 and constructed as indicated in Supplementary Table 5, Supplementary Table 6. Standard molecular cloning methods were used, and DNA assembly was performed using the NEBuilder HiFi mix (New England Biolabs). All oligos used in this study are listed in Supplementary Table 7. *B. bacteriovorus* strains were generated from the type strain HD100. *E. coli* strains used as prey were generated from MG1655. *E. coli* strains used for mating were generated from S17-λ*pir*. All plasmids were introduced in *B. bacteriovorus* by mating as described below. Scarless allelic replacements into the HD100 chromosome were performed using a strategy based on the two-step recombination a pK18mobsacB-derived suicide vector as in (74), screened by PCR and verified by DNA sequencing. Supplementary Table 8 contains information about all HD100 chromosomal regions used in this study. Protein fusions were confirmed by Western blot (Figure Supplement 10) and predation capacity of genetically-engineered strains was verified by killing curves (Figure Supplement 9), as described below.

### Routine culturing of *B. bacteriovorus* and *E. coli*

*E. coli* cells were routinely grown in LB medium except when otherwise stated. *B. bacteriovorus* strains were grown in DNB medium (Dilute Nutrient Broth, Becton, Dickinson and Company, supplemented with 2 mM CaCl_2_ and 3 mM MgCl_2_ salts) with *E. coli* as prey at 30°C with constant shaking as previously described (lysates) (75). Two-step revival of *B. bacteriovorus* from −80°C stocks was performed as in (75) except that only DNB medium was used. When appropriate, antibiotic-resistant *E. coli* strains were used as prey for overnight culturing of the corresponding antibiotics-resistant *B. bacteriovorus*. Kanamycin and gentamycin were used at 50 μg/ml and 10 μg/ml, respectively, both in liquid and solid media.

### Plasmid conjugation by mating

Mating was performed between *E. coli* S17-λ*pir* donor strain carrying the plasmid to be conjugated and the *B. bacteriovorus* receiver strain using a protocol modified from (74). Briefly, exponentially growing *E. coli* donor strains were harvested and washed twice in DNB medium before resuspension in 1:10 of the initial volume in DNB-salts. This donor suspension was mixed at equal volume with a fresh overnight lysate of a receiver HD100 strain. The mating mix was incubated for minimum 4 h at 30°C shaking before plating on selective medium using the double layer technique. Single plaques were isolated and transconjugants were confirmed by microscopy (when appropriate), PCR and sequencing.

### Labeling of *ori* and *ter*

The *B. bacteriovorus ori*_*Bb*_ (further named *ori*) has been located between the *dnaA* and *dnaN* genes (76) and the *ter_Bb_* region (*ter*) was identified by the 28-bp chromosome dimer resolution site *dif,* found between ORFs Bd2036 and Bd2038 (77). The *parS*_*PMT1*_ and *parS*_*P1*_ sequences were inserted near these loci by allelic replacement (between ORFs Bd3895 and Bd3896, and between ORFs Bd2052 and Bd2053, *i.e.* ~17 kbp and 12 kbp away from *ori*_*Bb*_ and *ter*_*Bb*_, respectively, Supplementary Table 10). We chose insertion sites in non-coding regions of ~60 nucleotides between 3’-ends of predicted ORFs to avoid interrupting transcription initiation signals. The insertion sites between Bd3895 and Bd3896, and between Bd2052 and Bd2053 are referred to as *ori* and *ter,* respectively, for simplicity; the strains in which *parS*_*PMT1*_ or *parS*_*P1*_ was inserted at those loci are referred to as *ori::parS*_*PMT1*_ and *ter::parS*_*P1*_, respectively.

### Live-cell imaging

*B. bacteriovorus* were first grown overnight with the appropriate *E. coli* prey and antibiotics if maintenance of a plasmid was required, then grown on wild-type MG1655 for at least one generation without antibiotic before the start of the imaging experiment. For snapshots of fresh AP *B. bacteriovorus*, cells were then spotted on 1.2% agarose pads prepared in DNB-salt media. For snapshots of *E. coli* strains, overnight cultures were diluted at least 1:500 and grown to exponential phase before being spotted on 1.2% agarose pads prepared in PBS or M9-salts buffer (supplemented with 0.2% glucose, 0.2% casamino acids and 1μg/ml thiamine, 2mM MgSO_4_ and 0.1mM CaCl_2_). For time-lapse or time-course imaging of synchronous predation cycles, MG1655 *E. coli* cells were grown in 2TYE medium to exponential phase (OD_600_ = 0.4-0.6), harvested at 2600 x g at RT for 5 minutes, washed twice and resuspended in DNB medium. Then, *E. coli* and *B. bacteriovorus* were mixed with a 1:3 to 1:5 volume ratio to allow most prey cells to be infected simultaneously. We consider the prey-predator mixing step as the time 0 in all our synchronous predation imaging experiments. Cells were either spotted directly on DNB-agarose pads for imaging, or left shaking at 30°C before imaging for the indicated durations. In time-lapse experiments, the same fields of view on the pad were imaged at regular interval times as indicated, with the enclosure temperature set to 28°C or 30°C. In time-course experiments, samples from the predation mixture were taken at regular interval times as indicated and directly spotted on agarose pads for snapshots. For nucleoid staining experiments, cells were incubated for 5 min prior imaging with DAPI (Sigma), SYTOX orange (Thermo Fisher) or Syto61 (Thermo Fisher) at a final concentration of 5 μg/ml, 500 nM and 200 nM, respectively. For flagellum staining, *B. bacteriovorus* AP cells were stained with the FM4-64 stain (Thermo Fisher) at a final concentration of 20 μg/ml and incubated in the dark for 2 min before detection or with CellBrite™ Fix 488 Membrane Dye (Biotium) at a final concentration of 10X (from a 1:1000 dilution) and incubated in the dark for 2 min before detection. For treatment with novobiocin, fresh AP *B. bacteriovorus* cells were mixed with prey as explained above, treated or not with 5 μg/ml novobiocin (Sigma) at the indicated times and before being immediately spotted on agarose pads containing 5 μg/ml novobiocin or not, respectively.

### EdU labelling of newly synthesized DNA

Newly synthesized DNA in *E. coli* and *B. bacteriovorus* cultures was labeled using the Click-iT EdU Alexa Fluor Imaging Kit (Invitrogen, Germany) as performed before with other bacteria (78,79). Briefly, 200 μl of *B. bacteriovorus* cells grown as indicated or *E. coli* cells grown exponentially in M9-glucose medium were incubated with ~12 μM 5-ethynyl-2′-deoxyuridine (EdU) for 5 and 15 minutes, respectively. Cells were fixed with 78% of ice-cold methanol to stop the reaction, washed in PBS (5000 x *g*, 4°C, 5 min), before membrane permeabilization in 100 μl PBS containing 0.5% Triton X-100 at room temperature for 30 minutes. Hereafter, the detergent was washed off twice with PBS. The pellet was resuspended in 40 μl of Click-iT reaction cocktail and incubated at room temperature covered from light for 30 minutes. The cells were collected, washed, resuspended in 40 μl of PBS, and when required treated with 5 μg/ml DAPI before imaging.

### Image acquisition

Phase contrast and fluorescence images were acquired on a Nikon Ti2-E fully-motorized inverted epifluorescence microscope (Nikon) equipped with CFI Plan Apochromat λ DM 100x 1.45/0.13 mm Ph3 oil objective (Nikon), a Sola SEII FISH illuminator (Lumencor), a Prime95B camera (Photometrics), a temperature-controlled light-protected enclosure (Okolab), and filter-cubes for DAPI, CFP, mCherry, YFP and GFP (Nikon). Multi-dimensional image acquisition was controlled by the NIS-Ar software (Nikon). Pixel size was 0.11 μm or 0.07 μm when using built-in 1X or 1.5X intermediate magnification, respectively. Identical LED illumination and exposure times were applied when imaging several strains and/or conditions in one experiment and were set to the minimum for time-lapse acquisitions to limit phototoxicity.

### Image processing

For figure preparation, images were processed with FIJI (80) keeping contrast and brightness settings identical for all regions of interest in each figure, except when otherwise stated. For Figures 5A, 6C, 7A and Figure Supplement 6B, D, denoising (Denoise.ai, Nikon) was applied on all phase contrast and fluorescence channels to improve the display of time-lapse images acquired with low exposure (which was required to preserve cell viability). Figures were assembled and annotated using Adobe Illustrator.

### Quantitative image analysis

Cell outlines were obtained with subpixel precision from phase contrast images for AP *B. bacteriovorus* cells, uninfected *E. coli* cells or entire bdelloplasts using the automated cellDetection tool in the open-source image segmentation and analysis software Oufti (81). For analysis of intracellular signal in *B. bacteriovorus* cells within bdelloplasts, predator cell outlines were manually added in Oufti using the same parameters as for AP cells detection. Fluorescent signals were added to cell meshes after background subtraction. Fluorescent foci and nucleoids were detected using the spotDetection and objectDetection modules in Oufti, respectively. We used the same optimized parameters for nucleoid detection on *B. bacteriovorus* and *E. coli* images (Supplementary Table 9). Parameters for spot detection were optimized for each dataset as described (81), except when analyzing biological replicates or for comparison between strains imaged under identical conditions, in which cases we used the same parameters optimized on the appropriate control set of images. Fluorescence-related analysis, nucleoids and spots-related information, as well as other properties of individual cells based on microscopy images were extracted from Oufti data and plotted using custom codes in Matlab (Mathworks), described in Supplementary Text 1. Demographs of relative fluorescence intensity in cells sorted by length were plotted as in (81,82). When needed, arrays of relative fluorescence were oriented based on the position of the maximal fluorescence intensity of the indicated signal in each cell half. Kymographs were obtained using the built-in kymograph function in Oufti (81). Colocalization was quantified manually for cells inside bdelloplasts. For Supplementary Figure 3, the development version of BactMAP (83) (github.com/vrrenske/BactMAP) was used to generate cell projections. Cells were oriented by shape and subsequently by their fluorescent focus, where cells without focus or with more than one focus were removed from the analysis. After this, the BactMAP function plotOverlay() was used to group cells by cell length into four equally-sized groups and plot the cell shape, DAPI shape and fluorescent spot localization of each cell, faceted by size group (Supplementary Text 1). Fluorescence intensity profiles along the centerline of individual cells (Figure 5B, Figure Supplement 3D, 5C) were obtained by plotting linescans from segmented lines drawn across each cell in FIJI (80).

### Statistical analyses

The sample sizes and number of repeats are included in the figure legends. Means, standard deviations and coefficients of variation (CV) were calculated in MATLAB (Mathworks) or Microsoft Excel. SuperPlots were generated in Microsoft Excel as described (84).

### Western blot analysis

Sample preparation for Western blot analysis was performed as in (85), starting from 3 ml in the case of cleared *B. bacteriovorus* lysates. Sample were loaded on NuPage Bis-Tris SDS precast polyacrylamide gels and ran at 190 V for 50 minutes in NuPAGE MES SDS running buffer. Western blotting was performed using standard procedures with the following primary antibodies: JL-8 monoclonal antibody (Takara) for GFP variants, YFP and CFP; polyclonal mCherry antibody (product # PA5-34974, Thermo Fisher) for mCherry. Signal from antibody binding was visualized by detecting chemiluminescence from the reaction of horseradish peroxidase with luminol and chemiluminescence was imaged with an Image Quant LAS 500 camera (GE Healthcare). Goat anti-mouse IgG-peroxidase antibody (Sigma) was used as a secondary antibody for JL-8. Goat anti-rabbit IgG-peroxidase antibody (Sigma) was used as a secondary antibody for mCherry. Antibodies were diluted following manufacturer’s recommendations. Figures were prepared using ImageJ and assembled and annotated in Adobe Illustrator.

### Killing curves

Killing curves assays (Figure Supplement 9) were performed after normalization of the *B. bacteriovorus* inoculum (using the SYBR Green assay described below). Equal amounts of predators from the same fresh cleared lysate were mixed with preys at a final OD_600_ of 0.1, and DNB medium was added to reach 150 μl per well in a transparent 96-well flat bottom plate. Technical triplicates were prepared in separate wells of the same plate in each experiment. The plate was shaken continuously (frequency 567 cpm (3mm)) at 30°C for 24 h in a Synergy H1m microplate reader (Biotek). Optical density measurements at 600 nm were taken every 20 minutes. Decrease of OD_600_ indicates prey lysis, as *B. bacteriovorus* cells do not affect absorbance. Features of the curves were extracted by fitting the data to an adaptation of the generalized sigmoid curve (Eq.1) using an R workflow based on a differential evolution algorithm. Briefly, a first rough fit is done using all the data to find an approximation of the sigmoid midpoint *s*. *s* is then used to estimate the sigmoid part of the data on which a second, more accurate fit is performed. Data were plotted using R programming language.

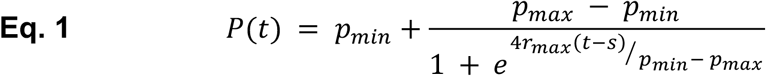

### SYBR Green normalization of predator cell density

For each cleared lysate of *B. bacteriovorus* to analyze, 198 μl/well were transferred into 3 wells of a black 96-well plate with transparent flat bottom (Sigma Aldrich). Then, protected from light, 2 μl of SYBR Green (LifeTechnologies) were added to each replicate to reach a volume of 200 μl per well. Plates without lid were shaken (double orbital, frequency 282 cpm (3mm)) in a Synergy H1m microplate reader (Biotek), for 15 min at 25°C before one end-point measurement of both OD_600_ and the SYBR Green fluorescence (490 nm excitation, 520 nm emission, gain 55). Based on a standard curve of OD_600_ relative to fluorescence values (obtained from serially diluted *E. coli* suspensions), the contribution of remaining *E. coli* in the lysates to the measured fluorescence was subtracted from the total SYBR Green fluorescence value in each well. The mean of the corrected fluorescence values from the 3 replicates is then used to compare the *B. bacteriovorus* density in different lysates and normalize them accordingly.

## Acknowledgements

We are grateful to Charles de Pierpont for excellent technical support, Michaël Deghelt for insightful discussions and critical review of the manuscript, Seung-Hyun Cho and Kilian Deconinck for advices on Western blots, Liz Sockett for providing the HD100 strain, Daniel Kadouri for plasmid pMQ414, Xavier De Bolle for *parS/*ParB constructs, Edouard Jurkevitch for plasmid pROBE-NT, Christine Jacobs-Wagner for sharing the unpublished strain CJW6321, Sarah Bigot for the construction of pBG18 and Christian Lesterlin for sharing the plasmid, Leendert Hamoen and Tanneke den Blaauwen for sharing plasmids pTNV143, pTNV162 and pTNV167, Pauline Leverrier and Boris Stojilković for critical reading of the draft. J.K. is a Research Fellow (Aspirant) of the F.R.S.-FNRS, T.L. is a FRIA grantee of the F.R.S.-FNRS, G.L. is a Research Associate (Chercheur Qualifié) of the F.R.S.-FNRS. Research in the Laloux lab is funded by the European Commission (ERC Starting Grant PREDATOR #802331) and the F.R.S.-FNRS (Incentive Grant for Scientific Research).

